# On Modeling the Macroecology of Baleen Whale Migration

**DOI:** 10.1101/009753

**Authors:** James R. Watson, Bruna Favetta, Charles Stock

## Abstract

Long distance migrations are well known to occur in many baleen whale species. Yet, a global synthesis of this information is lacking. Here, we study baleen whales as a group and at a global scale, first analyzing the grey and peer-reviewed literature for information on the location of baleen whale calving and feeding grounds around the world. This information was then combined with modeled-data produced from an Earth System Model to estimate the global distribution of baleen whale calving and feeding habitats. A simple network theoretic heuristic was then used to identify the shortest over-water path connecting habitats. These shortest paths map well to known major migration routes for a number of species, suggesting that migration has evolved primarily to minimize travel distances. Identifying distance minimizing routes globally, that have demonstrable consistency to known migration routes for certain baleen whale species, offers a useful baseline perspective on large-scale migration patterns, from which many perturbations can by judged. As an example, we used our modeled migration routes to identify regions of the ocean that are likely hotspots of whale ship-strikes. Such information is useful for developing global conservation and management priorities for baleen whales.

## Introduction

That baleen whales migrate is well known [1, 2]. One can observe them doing so from the numerous whale watching tours that are available around the world, and scientists have photographed and tagged individual whales, and tracked their travels [3–8]. From such data, and other sources of information, such as historical whaling charts [9], persistent long distance migration routes have been identified around the world, from the Gulf of California, the Hawaiian islands and the China Sea, to the Bering and Chuckchi Seas; from South Africa, Australia, New Zealand, Chile and Argentina, to the Southern Ocean and the Antarctic peninsula; from the Caribbean and the west coast of Africa, to the north Atlantic and Arctic. The list goes on. However, this information is rarely synthesized, with most studies focusing on migration patterns in specific regions and on specific species. Here, we make a global analysis of migration for baleen whales as a group, identifying the factors that likely determine migration patterns throughout the world’s oceans.

Annual migrations are commonly seen in baleen whales of different species, for example in hump-back [10] and gray whales [1], where individuals travel between warm low-latitude breeding grounds, and productive high-latitude feeding grounds [2]. Other baleen species are known to have more diffuse migration patterns, such as blue [5], sei [11] and fin [2] whales, but of those that make latitudinal migrations, they are thought to do so to secure better chances of survival for their calves in warmer low-latitude waters [2, 12], and feed on the abundance of copepods, euphausiids (krill) and small fishes at high latitudes in summer months [13, 14].

Low-to-high-to-low latitudinal migrations have been observed directly for many areas of the world’s oceans (see our Supplementary Information for a bibliography). Yet comparisons between regions are lacking, and a global synthesis of migration routes would develop our understanding of why certain baleen whale species, behave in such a (bioenergetically) expensive manner. Here, we develop a macroecological study, using network theoretic algorithms and output from an Earth System Model (ESM) to infer the location of migration pathways in the global ocean. ESMs have been developed primarily to explain the major biogeochemical cycles of the earth, ultimately giving us a predictive capacity for future climate states [15]. To make these calculations, earth-system scientists include descriptions of phytoplankton and (in some cases) zooplankton dynamics [16]. Here, we use a recent iteration of an ESM [17] that includes “large zooplankton”, which approximates the dynamics of copepods and euphausiids, as well as providing information useful for estimating the abundance of small fishes. This ESM data is interpolated to calving and feeding locations found from a literature review, and then a machine learning algorithm is used to define potential calving and feeding habitats globally. Major migration pathways between modeled calving and feeding habitats are then identified using a modified Dijkstra’s algorithm, which finds the shortest ocean-path between a given calving and feeding location.

Estimating the distribution of baleen whale calving and feeding habitats, and migration routes is not without uncertainty, for it is well known that baleen whale species differ in their use of the ocean. However, the global scale of our analysis offers a course grained view of species-specific information, and allows us to look at how baleen whales as a group, might occupy broad, basin-scale, regions of the global ocean. Furthermore, this large-scale perspective allows us to estimate the impact of humans on baleen whales, and we do so in the context of whale ship-strikes, thought to be a large source of mortality for migrating whales [18, 19]: using our modeled migration routes with cargo ship-track data, we infer hot-spots of potential ship-strikes globally.

## Methods

### The Earth System Model

The Carbon, Ocean Biogeochemistry and Lower Trophics (COBALT) marine ecosystem model provides the biogeochemical and planktonic food web information that we used to estimate potential whale migration routes [17]. COBALT is run as part of the Modular Ocean Model (MOM) version 4.1 [20], with 60 year simulations (1948-2008) forced by the Common Ocean-Ice Reference Experiment (CORE-II) data set [21]. The horizontal resolution of the simulation is 1*°* Latitude/Longitude, except along the equator where the resolution is refined to 1*/*3*°*, and the model uses 50 vertical layers, with a resolution of 10m over the top 200m.

The representation of planktonic food web dynamics within COBALT is based on body-size, and provides information relevant to our study of baleen whale migration. In particular, medium-sized zoo-plankton are parameterized to represent small and medium copepods (0.2 - 2*mm* in equivalent spherical diameter or ESD), and large-sized zooplankton are parameterized to represent large copepods and small euphasiids (2 - 20*mm* ESD). Furthermore, COBALT employs a density dependent mortality term for medium and large zooplankton. This represents the mortality of these zooplankton to higher-predators, namely small fishes. Although COBALT does not model fish explicitly [17], this implicit representation of small-fish feeding rates, has been used to model upper trophic levels explicitly [22] and for this application, provides information on the spatial distribution of a major source of food for certain baleen whale species.

### Baleen whale habitats

Data from the COBALT retrospective ocean-ice ecosystem simulation was used in a classification algorithm to identify likely baleen whale calving and feeding habitats globally. These data were chosen for their relevance to baleen whale habitats, and include: monthly COBALT temperature data averaged over the top 200m (*°C*), medium and large zooplankton (copepods and small euphausiids) biomass integrated over the top 200m (*g* carbon *m*^−2^; our analysis is based on relative difference in zooplankton abundances, so the exact units are unimportant), medium and large zooplankton mortality rates to small fishes integrated over the top 200m (*g* carbon *m*^−2^ *day*^−1^), depth (*km*). All data were taken for the period 1988-2008. Baleen whales generally calve during the winter, and feed during the summer, and hence these data were filtered for these times, producing time average austral winters and boreal summers (june, july, august), and boreal winters and austral summers (november, december, january). For illustrative purposes we show COBALT’s annual average sea-surface temperature (Fig. 1a) and average total abundance of medium and large zooplankton (Fig. 1b).

**Figure. 1.**
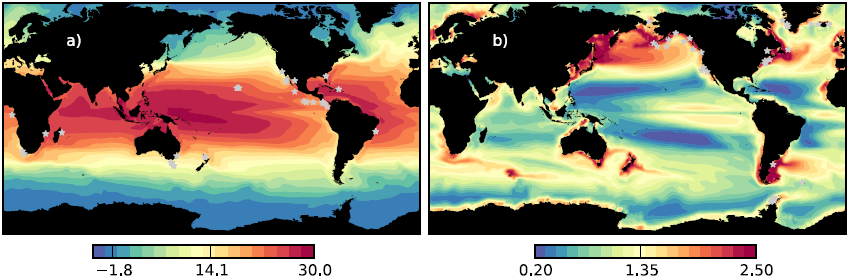
(a) Annual average sea surface temperature (*°C*) and (b) the annual average total abundance of medium and large zooplankton (*g* carbon *m*^−2^) from COBALT. Data are integrated over the top 200m, and baleen whale calving and feeding locations overlaid on Fig. 1a and b respectively (grey stars)

These time-averaged COBALT data fields were then linearly interpolated in space (the modeled data are on a 1*°* grid) to feeding and calving locations identified from a literature search: 33 and 36 feeding and calving sites respectively (Fig. 1). See the Supplementary Information for the bibliography of compiled literature, and the precise locations. Using this point-data, a machine learning algorithm - a bagged classification tree - was used to estimate the distribution of feeding and calving habitat at all other locations globally [23]. The algorithm was used on feeding and calving habitats separately. Taking feeding habitat as an example: the bagged classification tree required as input, a categorical response variable - a vector of 1s and 0s relating to those ocean locations where we identified the presence and absence of feeding respectively, from the literature. The other input is a matrix of predictor variables: the interpolated COBALT data at the feeding presence and absence locations. The output of the bagged classification tree, is another vector of 1s and 0s, relating to the classification of all ocean locations as either feeding habitat or not.

Absence locations were created by inspecting the distribution of zooplankton abundances, temperature, depth and irradiance across feeding and calving locations, and then using logical arguments to select ocean locations beyond the tails of these joint distributions. This essentially identifies areas of the ocean where calving and/or feeding does not happen. These are more accurately *pseudo-absence* locations because they are chosen, rather than identified [24], and for each run of any classification tree, 100 pseudo-absence points were selected randomly in this manner. With this information, the bagged classification tree then predicted the most likely habitat type (calving or feeding, or pseudo-absence) at every COBALT grid-cell, producing global maps of feeding and calving habitat.

Because the pseudo-absence points were chosen randomly, we repeated the classification process two hundred times, building a distribution of likely feeding, calving and pseudo-absence locations. Finally, because our concern is with the major migration routes, we isolated the most likely calving and feeding locations by choosing those areas with a 50% or greater probability of feeding or calving classification. Implicit to the bagged classification process is a k-fold cross-validation test of precision [25]. Here, part of the training data is left out and validated against, with the classification tree made using the remaining data. This process is repeated *k* times, producing an estimate of the expected classification error.

To assess the accuracy of the feeding and calving habitat classification trees, we compared its results with another set of information gained from the literature - we collected as much information as we could from peer and non-peer reviewed articles and numerous websites (representing the grey literature; see supplementary information) on the global distribution of calving and feeding habitats for a number of baleen whale species. The majority of this information was not geo-referenced in any detail, and as a result we had to visually overlay their maps, and estimate by-hand the distribution of baleen whale habitats and migration routes. This information was then compare qualitatively with the habitat distribution maps produced from the bagged-classficiation tree. This information was also used to divide the global feeding and calving habitat distributions, produced from the classification trees, into geographically consistent regions.

### Baleen whale migration pathways

Once global distributions of baleen whale calving and feeding habitats were made using the bagged-classificiation tree, we identified potential migration pathways using a modified Dijkstra’s algorithm [26]. This algorithm calculates the shortest-path between two nodes in a network. Here, we considered each COBALT grid-cell centroid a node in an ocean network. Then the great-circle distance (*km*) between each grid cell centroid and the centroid of every neighboring cardinal grid cell was calculated. These defined the edges in the ocean network. The result was a highly sparse matrix (or network): each row representing a grid-cell, with eight real-valued distances associated with its cardinal neighbors. Connections to land grid-cells were assigned an infinite distance. Dijkstra’s algorithm was then applied to this ocean network, identifying the shortest over-water or “ocean distance” between two given locations in the sea. However, because the network is constructed over a grid, unrealistic city-block paths were typical solutions. To solve this problem we added small amounts of white noise to the network edges, and applied Dijkstra’s algorithm a number of times (typically 1000), creating a set of candidate shortest paths. From the set of candidate shortest paths we calculated the most likely (or expected) shortest path between a given calving-feeding grid cell pair. Using this information, we identified the shortest path between calving and feeding regions. This approach is still a heuristic, and as a result the shortest paths are not exact solutions. An example shortest path is shown in Fig. 2. Here, numerous candidate paths are identified by the black dots, and the expected shortest path in red. The starting location is off Costa Rica, and the destination location is off Vancouver Island. The shortest path hugs the coast, and resembles the path taken by baleen whale species that migrate in this region (for example, blue whales [5]).

**Figure. 2.**
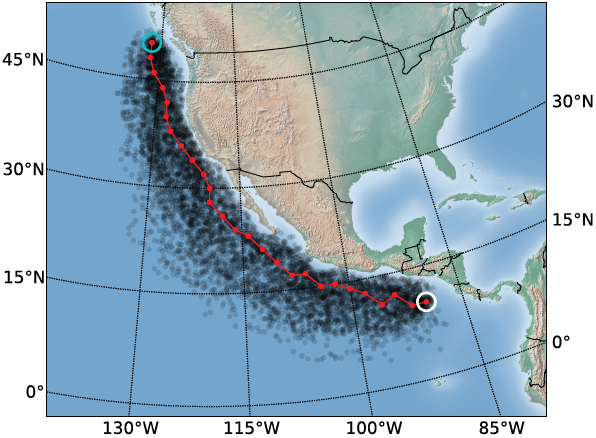
An outline of our route-finding algorithm. Black dots comprise locations along multiple candidate shortest paths, found using our modified Dijkstra’s algorithm, connecting the starting location (white circle) and terminus (blue circle). The red line is the expected shortest route, over all candidate shortest paths.

## Results

When compared qualitatively to the information gained from the literature (Fig. 3), the habitat classification algorithm identified realistic calving and feeding regions (Fig. 4; green and orange regions respectively; habitat regions are the same across all panels), and average cross-validation error for both calving and feeding trees was satisfyingly low (0.07 and 0.05 respectively after 50 trees; see Supplementary Information Fig. S1). Feeding habitats were identified mostly at high latitude: the Sea of Okhotsk, the Bering sea, the northwest Atlantic, the northeast Atlantic, Greenland, the North Sea, the Barents sea, the Crozet Islands, the south coast of Australia, southern New Zealand, the Patagonia Shelf and the whole of Antarctica. Calving habitats were found predominantly at in shallow, warm, low latitudes regions: Baja California, the off Costa Rica and Ecuador, off northwest Africa below the Cape Verde islands, the Gulf of Guinea, off the west-coast of Madagasca in the Mozambique channel, in the Arabian Sea, the Bay of Bengal, areas off the northwest and northeast coasts of Australia, and Fiji. These feeding and calving distributions map well to areas identified qualitatively from the peer-reviewed and grey literature (Fig. 3).

**Figure. 3.**
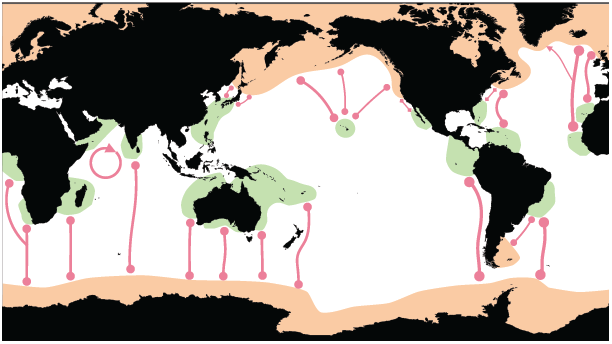
The distribution of baleen whale calving (green) and feeding (orange) habitats, and major migration routes, estimated qualitatively from the literature (peer and grey literature, as well as numerous website; see supplementary information). The major migration routes are generalizations of many specific regional examples, and are only meant to describe the major patterns of baleen whale migration at a global scale. Further, these routes are used to connect the feeding and calving regions, we do not attempt to describe whale movement within these habitats. The circular arrow in the Indian ocean identifies a resident group found there [36].

**Figure. 4.**
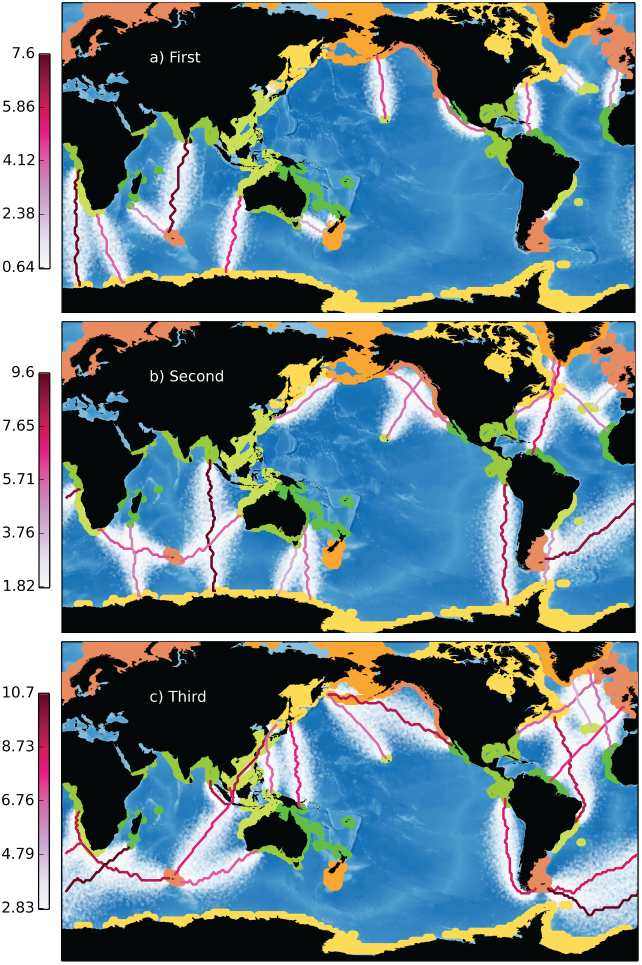
First (a), second (b) and third (c) shortest migration paths between all calving and feeding regions, identified using our modified Dijkstra’s algorithm. Calving regions are identified in green, the feeding regions in orange. The different shades of green and orange are used to simply differentiate the regions. Shortest paths are those that minimize ocean distance (colorbar: 1000**′***s km*). These paths are not exact solutions to the shortest-path problem, hence there are curves and bends to them. However, these do not impact the comparison of migration routes and distances between regions of the world’s oceans.

The partitioning of global habitat into geographically consistent regions, based on qualitative information gained from the literature, is shown in Fig. 4 (regions in different shades of green and orange). For example, the Hawaiian Islands, the Gulf of California, and the eastern tropical Pacific were all divided into separate calving regions. Similarly for the distribution of feeding habitat, areas such as Antarctica, the Bering Sea, and the North Sea are separated into distinct feeding regions. While agreement on a regional scale is good, there are a few noticeable calving regions not seen in the literature (Fig. 3). For example northern New Zealand and the Gulf of Mexico are classified as calving habitat, but there is little evidence for this [27]. Similarly for feeding grounds, the classification tree identified the southern half of New Zealand, but there was little evidence in the literature that this is a major feeding area for baleen whales. However, that is to say there is little evidence that *contemporary* whale distributions cover these areas [27]. Both are interesting as they constitute potential feeding and calving regions for right whales, species which have been essentially exterminated and may show significant range contraction. There is evidence that Southern New Zealand was a feeding ground and Northern New Zealand was a calving area for Southern Right Whales before they were almost wiped out [28]. Similarly, the Gulf of Mexico may have been an extension of the Florida East Coast calving area for Northern Right Whales, with a few individuals spotted in the GOM in the past several decades. Considering there are only about 400 Northern Right Whales currently, it seem highly plausible that the GOM/Caribbean was an important calving area for them when they numbered in the 10’s of thousands [29].

Globally, the first, second and third shortest paths that minimize over-water distance are shown in Fig. 4a, b & c respectively. The maximum, minimum and mean path lengths (*km*), for these three sets are: first - 7291, 696, 3037; second - 9469, 2091, 5014; third - 10828, 2414, 6948. Beyond these simple statistics, what is evident are the realistic migration routes. These shortest paths between every calving region and their three nearest feeding regions describe all migration routes identified in the grey literature (Fig. 3). For example, from Hawaii, the three nearest feeding regions are the Sea of Okhotsk, the Bering Sea, and the waters off the northwest US and Canada. From the southeast Caribbean, shortest migration routes connect to the waters of the northeast US and Canada, to Greenland, and to the North Sea. In the southern hemisphere, many calving locations connect to the Southern Ocean and Antarctica, traversing many degrees of latitude, for little change in longitude. There are however shortest paths that traverse several degrees on longitude. These are most commonly the third shortest paths, for example connecting Madagascar and northwest Australia, with the Crozet Islands (Fig. 4c).

## Discussion

Baleen whale feeding and calving regions were identified, using peer-reviewed and grey literature information on calving and feeding locations, within a bagged-classification tree. Then, using a modified Dijkstra’s algorithm, we identified paths that minimized the ocean distance between feeding and calving regions. Both the global distribution of feeding and calving habitats and the shortest paths connecting them, map well to the known distributions of these habitats and migration routes for certain species. Our results suggest that baleen whale migration has evolved to minimize ocean-distances between calving and feeding habitats. Other factors, such as predator avoidance (e.g. from killer whales) are likely to play a role too in shaping where baleen whales go, but at smaller spatial scales [12].

This analysis is global in scale, and baleen whales were analyzed as a group. This latter step places constraints on our analysis, as it is known that migration patterns, and behavior in general, can differ greatly between whale-species. This is reflected in the specificity of most whale-movement data, which is typically collected for particular individuals and groups of certain whale species, in specific regions, where focused observational and tagging efforts can give fine-scale knowledge of movements [5, 8]. The global-scale analysis presented herein lacks the detail of regional, species specific studies, however, it provides a coarser global perspective on large-scale movements of baleen whales and the mechanisms underlying them. Indeed, the large scale of our analysis does not preclude species-specific use: different baleen whale species show common reliance on broad ocean areas (at scales: *>* 1000*s km*) citeGregr2001, and hence our modeled baleen whale habitat maps and migration routes may capture patterns that are general across species. Furthermore, simply having a global atlas of possible migration routes allows us to gauge how humans may have impacted whale migrations around the world.

As an example, we use our global atlas of baleen whale migration to identify hot-spots of potential ship strikes. Ship strikes are thought to be a large source of mortality for some whale groups [18, 19]. But like assessments of migration, most analyses of ship strikes are at regional scales. In contrast, our work here can help identify global patterns of ship strike potential. Using information from the PASTA MARE project [30], we interpolated normalized shipping density (a score between [0,1]; see Fig. S2 for a detailed description) to locations along the major whale migration pathways, resulting in the identification of potential ship-strike hot-spots (Fig. 5). Japanese waters emerge as having the highest strike risk for a given whale migrating in this area. Secondary hot-spots occur in Indonesian waters, off South Africa, off Puerto Rico and off New Foundland, Canada. Although this brief analysis looks only at ship strike potential along migration routes, ignoring strikes in calving and feeding regions, this information can be useful for developing global conservation and management priorities. Two important caveats to this analysis is that first, these strike risk estimates ignore any regional-scale measures to reduce risk, for example as has been successfully done for the North Atlantic Right Whale [31]. Second, we assume a *pristine* ocean, for we do not account for the effect of historic whaling on contemporary whale densities. Thus, inferring ship-strike potential in this way is likely inaccurate due to the loss of migration routes/populations from whaling [9]. Indeed, a worthwhile effort is the comparison of contemporary migration routes with our estimates of migration in a pristine ocean. Through this, one can identify (or at least hypothesize for) the loss of particular migration routes.

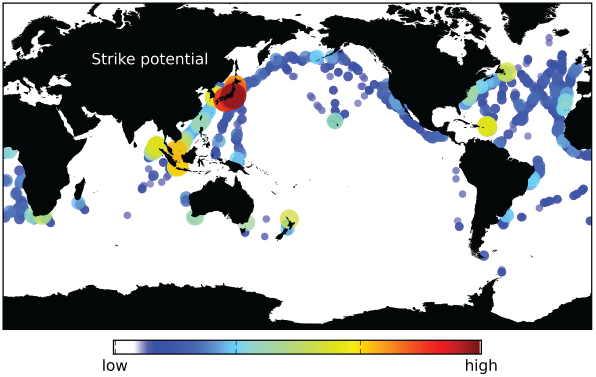
Baleen whale strike potential. Along with color, the size of the dots denotes whale strike potential: the larger the higher potential. See Fig. S2 for a description of the shipping data.

Looking to the past is important, but equally so is to understand how humans may continue to impact whale migration in the future. Our feeding and calving habitat maps, and migration routes were trained on Earth System Model data, averaged over the period 1988-2008. As a result our modeled distributions of feeding and calving habitats are specific to this period. It is possible too, to use our techniques to project how habitat distributions and migration routes might change in the future. The Earth System Model data used in this analysis, as well as looking back in time in hindcast simulations, also look forward in time and project the distribution of oceanographic variables such as temperature and the abundance of zooplankton [15, 17]. Thus, with this information it is possible, in future work, to estimate how baleen whale migration feeding and calving habitats might move under different future climate scenarios.

To improve the accuracy of these future projections, especially the estimates of migration routes, there are several modifications that could be made to our *basic* approach. We say basic because our migration route-finding algorithm does not include any biological information. Hence, we cannot answer the question of what condition whale individuals might be in, for example, if calving and feeding habitats move further apart in the future, or if their quality (e.g. the abundance of large zooplankton or krill) might diminish. It is critical to do so because individual condition can have a highly non-linear relationship with travel-distance/time [32], and slight increases in the shortest migration path distance, could be detrimental to whale health, rendering certain migration routes impossible. The experience along a migration route is important too. For example, it is expected that areas of the surface ocean are likely to warm in the coming decades. This may incur a significant bioenergetic cost on migrating whales, as biomass specific respiration rates are positively related to temperature [33]. In other words, baleen whales are likely to expend more energy on their annual migrations in the future, and the consequences of this change are unknown.

In addition, there are several algorithmic modifications to our approach that can be made. First, we have only identified paths that minimize ocean distance. It is possible too, to find paths between two locations is the sea that minimize travel-time. We have done so in our context (Fig. S3), assuming that baleen whales swim with a constant speed of 80 *km day*^−1^ [5], and instead of distances as network edges (see methods) we assign travel times, accounting for ocean currents. Applying Dijkstra’s algorithm to this network creates solutions to what is known as the *Zermelo navigation problem*, that is, finding the path that minimizes travel time between two locations in the sea [34]. The resulting migration routes are shown in Fig. S3, and we find little difference between these and the distance minimizing routes. This is due to the fast swimming speeds of baleen whales. However, solutions to Zermelo’s problem for other species that swim more slowly, for example for marine turtles [34], have identified multiple possible migration routes, depending on the movement rule employed. This brief analysis reveals the numerous ways in which potential migration routes can be identified, and how this information can be used to develop hypotheses about the optimal movement strategies that marine species have evolved.

In summary, we estimated the distribution of baleen whale feeding and calving habitats globally, and the migration routes connecting them. We have shown evidence that baleen whales, as a group, have migration routes that minimize distance (and/or travel time), and we have also discussed that the main utility of our macroecological analyses is in the generation of hypotheses and questions. For example, do baleen whales minimize travel distance or time? Answers to these questions may come from evolutionary agent-based simulations that have been employed in idealized settings to answer similar questions [35]. Indeed, combining these tools with Earth System Model data, will better position us to understand where baleen whales migrate to and from, how they do it, and why.

## Acknowledgments

We would like to thank Andre Boustany, Jorge Sarmiento, Simon Levin, Malin Pinsky, Rebecca Asch and Jodi Elliott for their help in preparing this work. JW was part funded by the Nippon Foundation (NF) Nereus Program. JW was also funded by the NSF Coupled Natural-Human Systems grant GEO-1211972. BF was funded by Princeton University’s Grand Challenges program.

## Supplementary Information 1

### Literature Review

We collected information on the distribution of baleen whale calving and feeding habitats, and their migration routes. Quantitative - geo-referenced - information was obtained from peer-reviewed literature and is listed in Table A1, with corresponding articles in the subsequent bibliography. Qualitative information - non geo-referenced - from the grey-literature: websites and non-peer reviewed publications are listed at the end.

**Table 1.** Habitat type, location, species and reference number for literature identifying baleen whale calving and/or feeding locations. References are listed in the bibliography below.

| Habitat type | Latitude | Longitude | Species | Reference |
| --- | --- | --- | --- | --- |
| Calving | 5 | -80 | Humpback | 1 |
| Calving | 18 | -108 | Humpback | 2 |
| Calving | 20 | -150 | Humpback | 2 |
| Calving | 20 | -160 | Humpback | 2 |
| Calving | 20 | -156 | Humpback | 11 |
| Calving | -16 | -38 | Humpback | 13 |
| Calving | -32 | 17 | Right | 14 |
| Calving | -18 | 37 | Humpback | 15 |
| Calving | -2 | 8 | Humpback | 16 |
| Calving | -17 | 51 | Humpback | 16 |
| Calving | 9 | -83 | Humpback | 17 |
| Calving | 9 | -84 | Humpback | 17 |
| Calving | 9 | -84 | Humpback | 17 |
| Calving | 8 | -82 | Humpback | 17 |
| Calving | 9 | -84 | Humpback | 17 |
| Calving | -41 | 145 | Right | 18 |
| Calving | -36 | 175 | Right | 18 |
| Calving | -36 | 150 | Right | 18 |
| Calving | -34 | 18 | Right | 18 |
| Calving | -43 | 147 | Right | 18 |
| Calving | -37 | 174 | Right | 18 |
| Calving | -41 | 145 | Right | 18 |
| Calving | -43 | 148 | Right | 18 |
| Calving | 9 | -92 | Blue | 19 |
| Calving | 9 | -98 | Blue | 19 |
| Calving | 28 | -115 | Blue | 20 |
| Calving | 26 | -115 | Blue | 20 |
| Calving | 34 | -119 | Blue | 20 |
| Calving | 10 | -100 | Blue | 20 |
| Calving | 31 | -81 | Right | 20 |
| Calving | -42 | -63 | Right | 23 |
| Calving | -16 | -38 | Humpback | 24 |
| Feeding | 35 | -125 | Humpback | 1 |
| Feeding | 60 | -147 | Humpback | 2 |
| Feeding | 55 | -140 | Humpback | 2 |
| Feeding | 40 | -70 | Humpback | 3 |
| Feeding | 50 | -50 | Humpback | 3 |
| Feeding | 65 | -15 | Humpback | 3 |
| Feeding | 67 | -55 | Humpback | 4 |
| Feeding | 64 | -51 | Humpback | 5 |
| Feeding | 65 | -30 | mix (fin,blue, sei, humpback) | 6 |
| Feeding | 65 | -51 | mix (fin,blue, sei, humpback) | 7 |
| Feeding | 64 | -50 | mix (fin,blue, sei, humpback) | 8 |
| Feeding | 50 | -50 | mix (humpback, fin, minke) | 9 |
| Feeding | -64 | -63 | Humpback | 10 |
| Feeding | -64 | -62 | Humpback | 12 |
| Feeding | -64 | -63 | Humpback | 17 |
| Feeding | -65 | -65 | Humpback | 17 |
| Feeding | -65 | -63 | Humpback | 17 |
| Feeding | -65 | -64 | Humpback | 17 |
| Feeding | -65 | -62 | Humpback | 17 |
| Feeding | 41 | -69 | Right | 21 |
| Feeding | 65 | -24 | Minke | 22 |
| Feeding | -54 | -38 | Right | 23 |
| Feeding | -63 | -62 | Humpback | 12 |
| Feeding | 50 | -132 | Gray | 25 |
| Feeding | 67 | -170 | Gray | 26 |
| Feeding | 49 | -68 | mix(fin, minke) | 27 |
| Feeding | 42 | -70 | Humpback | 28 |
| Feeding | 43 | -66 | Humpback | 28 |
| Feeding | 34 | -120 | Humpback | 28 |
| Feeding | 37 | -123 | Humpback | 28 |
| Feeding | 48 | -125 | Humpback | 28 |
| Feeding | 57 | -135 | Humpback | 28 |
| Feeding | 56 | -153 | Humpback | 28 |
| Feeding | 52 | -155 | Humpback | 28 |
| Feeding | 52 | -163 | Humpback | 28 |
| Feeding | 54 | -166 | Humpback | 28 |

## Whale Migration Websites

w3.shorecrest.org,

www.whales.org.au

www.ecy.wa.gov

www.learner.org,

www.marinebio.net

www.whalesbc.com

www.ocean-institute.org

www.whoi.edu

www.wildhawaii.org

www.mersea.com

www.nature.nps.gov

www.kids.britannica.com

www.library.arcticportal.org

www.science.kqed.org

www.bajainsider.com

www.csiwhalesalive.org

http://nationalgeographic.pny.com/Mapshome.aspx?Category=Maps

www.coastalstudies.org

www.phys.org

www.scrippsblogs.ucsd.edu,

www.buzzle.com

www.whalemuseum.is

www.beforeitsgone.com.au

http://www.apex-environmental.com/

http://www.ecoflores.org/en/marine+conservation/supporting+foreign+ngo/apex/

## Supplementary Figures

**Figure S1:**
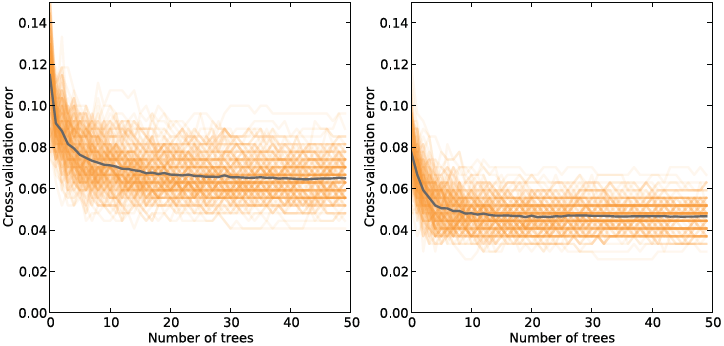
(a). Habitat modeling cross-validation error for calving (left) and feeding (right) habitats. The classification tree algorithm is applied multiple times (orange lines), as pseudo-absence locations are chosen at random. This results in an expected cross-validation error (grey line).

**Figure S2:**
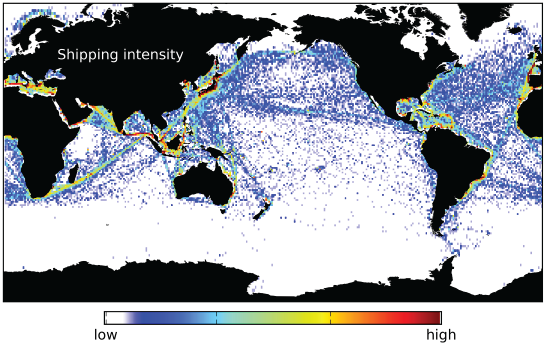
Ship density from the PASTA MARE project. This is defined as the average number of vessels within a grid cell, based on 10 global Satellite-Automatic Identification System (S-AIS) scenes. Each global S-AIS scene retains one position report per vessel within a time frame of 8 days. The data used covers the period from 1 January 2010 – 31 March 2010. We log-transformed this data then normalized it to [0,1], before interpolating to locations along the major migration path-ways, inferred from our ocean-distance minimizing algorithm. For more information on this data see https://webgate.ec.europa.eu/maritimeforum/content/1603

**Figure S3:**
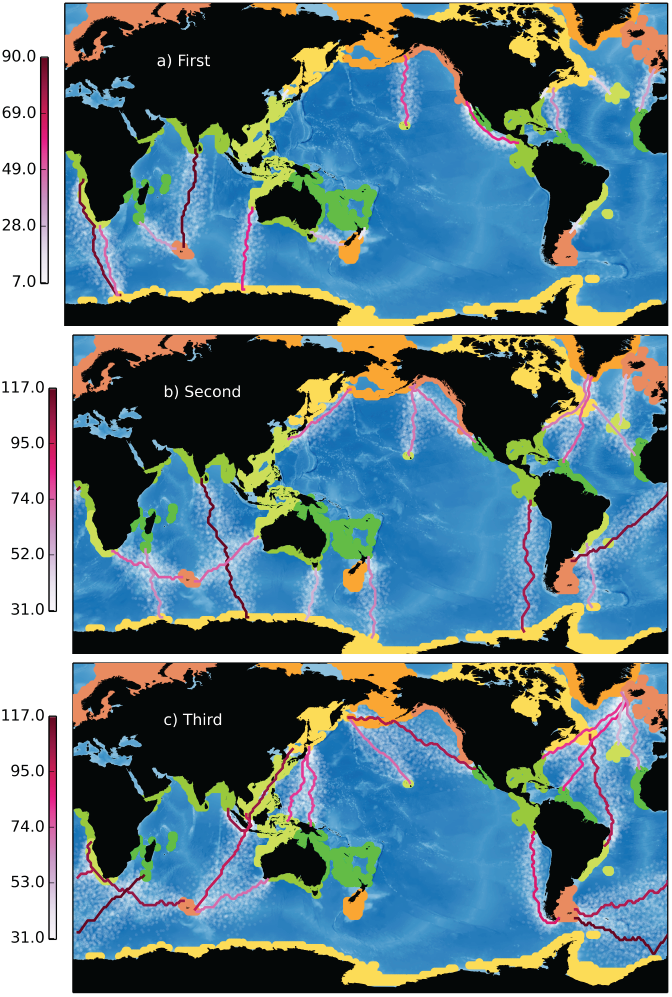
First (a), second (b) and third (c) shortest migration paths between all calving and feeding regions. Calving regions are identified in green, the feeding regions in orange. The different shades of green and orange are used to simply differentiate the regions. Shortest paths are those that minimize travel time (colorbar: *days*); assuming a constant swimming speed of 80 *km day*^−1^

## References

1. Pike GC (1962) Migration and feeding of the gray whale (Eschrichtius gibbosus). Journal of the Fisheries Board of Canada.

2. Corkeron P, Connor R (1999) Why do baleen whales migrate? Marine Mammal Science 15: 1228–1245.

3. Calambokidis J, Barlow J (2004) Abundance of blue and humpback whales in the eastern north pacific estimated by capture-recapture and line-transect methods. Marine Mammal Science 20: 63–85.

4. Dalla Rosa L, Secchi ER, Maia YG, Zerbini AN, Heide-Jorgensen MP (2008) Movements of satellite-monitored humpback whales on their feeding ground along the Antarctic Peninsula. Polar Biology 31: 771–781.

5. Bailey H, Mate BR, Palacios DM, Irvine L, Bograd SJ, et al. (2010) Behavioural estimation of blue whale movements in the Northeast Pacific from state-space model analysis of satellite tracks. Endangered Species Research 10: 93–106.

6. Boye TK, Simon M, Madsen PT (2010) Habitat use of humpback whales in Godthaabsfjord, West Greenland, with implications for commercial exploitation. Journal of the Marine Biological Association of the United Kingdom 90: 1529–1538.

7. Fretwell PT, Staniland IJ, Forcada J (2014) Whales from space: counting southern right whales by satellite. PLoS one 9: e88655.

8. Rosenbaum HC, Maxwell SM, Kershaw F, Mate B (2014) Long-Range Movement of Humpback Whales and Their Overlap with Anthropogenic Activity in the South Atlantic Ocean. Conservation biology: the journal of the Society for Conservation Biology 28: 604–615.

9. Smith TD, Reeves RR, Josephson EA, Lund JN (2012) Spatial and Seasonal Distribution of American Whaling and Whales in the Age of Sail. PLoS ONE 7: e34905.

10. Rasmussen K, Palacios DM, Calambokidis J, Saborío M.T, Dalla Rosa L, et al. (2007) Southern Hemisphere humpback whales wintering of Central America: insights from water temperature into the longest mammalian migration. Biology Letters 3: 302–305.

11. Mizroch S, Rice D, Breiwick J (1984) The sei whale, Balaenoptera borealis. Marine Fisheries Review 46: 25–29.

12. Jefferson T.A, Stacey P.J, Baird R.W (1991) A review of Killer Whale interactions with other marine mammals: predation to coexistence. Mammal Review 21: 151–180.

13. Kann L.M, Wishner K (1995) Spatial and temporal patterns of zooplankton on baleen whale feeding grounds in the southern Gulf of Maine. Journal of Plankton Research 17: 235–262.

14. Branch T, Staford K.M, Palacios D.M, Allison C, Bannister J.L, et al. (2007) Past and present distribution, densities and movements of blue whales Balaenoptera musculus in the Southern Hemisphere and northern Indian Ocean. Mammal Review 37: 116–175.

15. Dunne J.P, John J.G, Adcroft A.J, Griffies S.M, Hallberg R.W, et al. (2012) GFDL’s ESM2 Global Coupled Climate–Carbon Earth System Models. Part I: Physical Formulation and Baseline Simulation Characteristics. Journal of Climate 25: 6646–6665.

16. Stock C.A, Alexander M.A, Bond N.A, Brander K.M, Cheung WWL, et al. (2011) On the use of IPCC-class models to assess the impact of climate on Living Marine Resources. Progress In Oceanography 88: 1–27.

17. Stock C.A, Dunne J.P, John J.G (2014) Global-scale carbon and energy flows through the marine planktonic food web: An analysis with a coupled physical–biological model. Progress In Oceanography 120: 1–28.

18. Knowlton A.R, Kraus S.D (2001) Mortality and serious injury of northern right whales (Eubalaena glacialis) in the western North Atlantic Ocean. Journal of Cetacean Research and Management (Special Issue) 2: 193–208.

19. Jensen A.S, Silber G.K, Calambokidis J (2003) Large whale ship strike database. NOAA Technical Memorandum NMFS-OPR-: 37.

20. Griffies S.M (2012) Elements of the Modular Ocean Model (MOM): 2012 release. GFDL Ocean Group Technical Report No 7: 1–631.

21. Large W.G, Yeager S.G (2008) The global climatology of an interannually varying air–sea flux data set. Climate Dynamics 33: 341–364.

22. Watson J, Favetta B, CAS (In press) Exploring the role of movement in determining the global distribution of marine biomass using a coupled hydrodynamic –size-based ecosystem model. Progress in Oceanography.

23. Prasad A.M, Iverson L.R, Liaw A (2006) Newer Classification and Regression Tree Techniques: Bagging and Random Forests for Ecological Prediction. Ecosystems 9: 181–199.

24. Engler R, Guisan A, Rechsteiner L (2004) An improved approach for predicting the distribution of rare and endangered species from occurrence and pseudo?absence data. Journal of Applied Ecology 41: 263–274.

25. Elith J, Leathwick J.R, Hastie T (2008) A working guide to boosted regression trees. The Journal of Animal Ecology 77: 802–813.

26. Golden B (1976) Shortest-path algorithms: A comparison. Operations Research: 1164–1168.

27. Constantine R, Russell K (2007) Photo-identification of humpback whales in New Zealand waters and their migratory connections to breeding grounds of Oceania. Marine Mammal Science 23: 715–720.

28. Baker C.S, Patenaude N.J, Bannister J.L, Robins J, Kato H (1999). Distribution and diversity of mtDNA lineages among southern right whales (Eubalaena australis) from Australia and New Zealand. DOI:10.1007/s002270050519.

29. Ward-Geiger L, Knowlton A.R, Amos A.F, Pitchford T.D, MaseGuthrie B, et al. (2011). Recent Sightings of the North Atlantic Right Whale in the Gulf of Mexico.

30. Eiden G, Martinsen T (2010) Maritime traffic density - results of PASTA MARE project. Technical Note 41 Vessel Density Mapping Issue 4 Preparatory Action for Assessment of the Capacity of Spaceborne Automatic Identification System Receivers to Support EU Maritime Policy.

31. Conn P.B, Silber G.K (2013) Vessel speed restrictions reduce risk of collision-related mortality for North Atlantic right whales. Ecosphere 4: 1–15.

32. Wiedenmann J, Cresswell K.A, Goldbogen J, Potvin J, Mangel M (2011) Exploring the effects of reductions in krill biomass in the Southern Ocean on blue whales using a state-dependent foraging model. Ecological Modelling 222: 3366–3379.

33. Brown J, Gillooly J, Allen A.P, Savage V.M, West G (2004) Toward a metabolic theory of ecology. Ecology 85: 1771–1789.

34. Hays G.C, Christensen A, Fossette S, Schofield G, Talbot J, et al. (2014) Route optimisation and solving Zermelo’s navigation problem during long distance migration in cross flows. Ecology Letters 17: 137–143.

35. Shaw A.K, Couzin I.D (2013) Migration or Residency? The Evolution of Movement Behavior and Information Usage in Seasonal Environments. The American Naturalist 181: 114–124.

36. Mikhalev Y.A (1997) Humpback whales Megaptera novaeangliae in the Arabian Sea. Marine Ecology Progress Series 149: 13–21.

## Biliography

1. Calmbokidis, J., et al. (2000). Migratory destinations of humpback whales that feed of California, Oregon and Washington. Marine Ecology Progress Series, 192, 295–304.

2. Calambokidis, J., et al. (2001). Movements and population structure of humpback whales in the north pacific. Marine Mammal Science, 17(4), 769–794.

3. Stevick, P. T., et al. (2003). Segregation of migration by feeding ground origin in North Atlantic humpback whales (Megaptera novaeangliae). Journal of Zoology, 259(3), 231–237. DOI:10.1017/S0952836902003151

4. Simon, Malene Juul. “The sounds of whales and their food: Baleen whales, their foraging behaviour, ecology and habitat use in an arctic habitat.” PhD dissertation., Aarhus Universitet Aarhus University, Department of Biological Sciences, Zoophysiology, 2010.

5. Boye, T. K., et al. (2010). Habitat use of humpback whales in Godthaabsfjord, West Greenland, with implications for commercial exploitation. Journal of the Marine Biological Association of the United Kingdom, 90(08), 1529–1538.

6. Heide-Jorgensen, M. P., & Simon, M. J. (2007). Estimates of large whale abundance in Greenlandic waters from a ship-based survey in 2005. Journal of Cetacean Research and Management, 9(2), 95–104.

7. Heide-Jorgensen, M. P., & Laidre, K. L. (2013). Surfacing time, availability bias, and abundance of humpback whales in West Greenland. International Whaling Commission, 65a(AWMP1), 1–13.

8. Heide-Jorgensen, M. P., et al. (2007). Increasing abundance of bowhead whales in West Greenland. Biology Letters, 3(5), 577–580. DOI:10.1098/rsbl.2007.0310

9. Piatt, J. F., et al. (1989). Baleen whales and their prey in a coastal environment. Canadian Journal of Zoology, 67(6), 1523–1530.

10. Friedlaender, A. S., et al. (2008). The effects of prey demography on humpback whale (Megaptera novaeangliae) abundance around Anvers Island, Antarctica. Polar Biology, 31(10), 1217–1224.

11. Cartwright, R., & Sullivan, M. (2009). Behavioral ontogeny in humpback whale (Megaptera novaeangliae) calves during their residence in Hawaiian waters. Marine Mammal Science, 25(3), 659–680.

12. Dalla Rosa, L., et al. (2008). Movements of satellite-monitored humpback whales on their feeding ground along the Antarctic Peninsula. Polar Biology, 31(7), 771–781.

13. Lunardi, D. G., et al. (2010). Behavioural strategies in humpback whales, Megaptera novaeangliae, in a coastal region of Brazil. Journal of the Marine Biological Association of the United Kingdom, 90(08), 1693–1699.

14. Fortune, S. (2012). North Atlantic right whale growth and energetics. Masters Thesis. Department of Zoology. The University of British Columbia.

15. Pomilla, C., & Rosenbaum, H. C. (2005). Against the current: an inter-oceanic whale migration event. Biology Letters, 1(4), 476–479.

16. Pomilla, C., & Rosenbaum, H. C. (2006). Estimates of relatedness in groups of humpback whales (Megaptera novaeangliae) on two wintering grounds of the Southern Hemisphere. Molecular Ecology, 15(9), 2541–2555.

17. Rasmussen, K., Palacios, D. M., Calambokidis, J., Saboro, M. T., Dalla Rosa, L., Secchi, E. R., et al. (2007). Southern Hemisphere humpback whales wintering of Central America: insights from water temperature into the longest mammalian migration. Biology Letters, 3(3), 302–305.

18. Kemper, C. M. (2002). Distribution of the pygmy right whale, Caperea marginata, in the Australasian region. Marine Mammal Science, 18(1), 99–111.

19. Calambokidis, J., & Barlow, J. (2004). Abundance of blue and humpback whales in the eastern north pacific estimated by capture-recapture and line-transect methods. Marine Mammal Science, 20(1), 63–85.

20. Mate, B. R., Lagerquist, B. A., & Calambokidis, J. (1999). movements of north pacific blue whales during the feeding season of southern california and their southern fall migration. Marine Mammal Science, 15(4), 1246–1257.

21. Kann, L. M., & Wishner, K. (1995). Spatial and temporal patterns of zooplankton on baleen whale feeding grounds in the southern Gulf of Maine. Journal of Plankton Research, 17(2), 235–262.

22. Christiansen, F., Rasmussen, M., & Lusseau, D. (2013). Whale watching disrupts feeding activities of minke whales on a feeding ground. Marine Ecology Progress Series, 478, 239–251.

23. Valenzuela, L. O., Sironi, M., Rowntree, V. J., & Seger, J. (2009). Isotopic and genetic evidence for culturally inherited site fidelity to feeding grounds in southern right whales (Eubalaena australis). Molecular Ecology, 18(5), 782–791.

24. Danilewicz, D., Tavares, M., Moreno, I. B., Ott, P. H., & Trigo, C. C. (2009). Evidence of feeding by the humpback whale (Megaptera novaeangliae) in mid-latitude waters of the western South Atlantic. Marine Biodiversity Records, 2, e88.

25. Pike, G. C. (1962). Migration and feeding of the gray whale (Eschrichtius gibbosus). Journal of the Fisheries Board of Canada.

26. Sobolevskii, E. I., Yakovlev, Y. M., & Kusakin, O. G. (2000). Some data on macrobenthos composition on feeding grounds of the gray whale (Eschrichtius gibbosus Erxl., 1777) on the northeastern Sakhalin shelf. Russian Journal of Ecology, 31(2), 126–128.

27. Simard, Y., Lavoie, D., & Saucier, F. J. (2002). Channel head dynamics: capelin (Mallotus villosus) aggregation in the tidally driven upwelling system of the Saguenay - St. Lawrence Marine Park’s whale feeding ground. Canadian Journal of Fisheries and Aquatic Sciences, 59(2), 197–210. DOI:10.1139/f01-210

28. Elfes, C. T., Vanblaricom, G. R., Boyd, D., Calambokidis, J., Clapham, P. J., Pearce, R. W., et al. (2010). Geographic variation of persistent organic pollutant levels in humpback whale (Megaptera novaeangliae) feeding areas of the North Pacific and North Atlantic. Environmental Toxicology and Chemistry, 29(4), 824–834. DOI:10.1002/etc.110

